# An atlas of protein-protein interactions across mammalian tissues

**DOI:** 10.1101/351247

**Authors:** Michael A. Skinnider, Nichollas E. Scott, Anna Prudova, Nikolay Stoynov, R. Greg Stacey, Joerg Gsponer, Leonard J. Foster

## Abstract

Cellular processes arise from the dynamic organization of proteins in networks of physical interactions. Mapping the complete network of biologically relevant protein-protein interactions, the interactome, has therefore been a central objective of high-throughput biology. Yet, because widely used methods for high-throughput interaction discovery rely on heterologous expression or genetically manipulated cell lines, the dynamics of protein interactions across physiological contexts are poorly understood. Here, we use a quantitative proteomic approach combining protein correlation profiling with stable isotope labelling of mammals (PCP SILAM) to map the interactomes of seven mouse tissues. The resulting maps provide the first proteome-scale survey of interactome dynamics across mammalian tissues, revealing over 27,000 unique interactions with an accuracy comparable to the highest-quality human screens. We identify systematic suppression of cross-talk between the evolutionarily ancient housekeeping interactome and younger, tissue-specific modules. Rewiring of protein interactions across tissues is widespread, and is poorly predicted by gene expression or coexpression. Rewired proteins are tightly regulated by multiple cellular mechanisms and implicated in disease. Our study opens up new avenues to uncover regulatory mechanisms that shape *in vivo* interactome responses to physiological and pathophysiological stimuli in mammalian systems.

## Introduction

Cellular functions are mediated by the dynamic association of individual proteins into complexes, signalling pathways, and other macromolecular assemblies. The biological functions of many proteins depend on specific physical interactions with other proteins, and disruption of these interactions can result in disease (Sahni et al., 2015). Defining the complete map of functional protein-protein interactions in a given organism (the interactome) has therefore been a longstanding goal of the post-genomic era, with a view to better understanding protein function, cellular processes, and ultimately the relationship between genotype and phenotype (Vidal et al., 2011). To this end, high-throughput methods have been developed to map interactomes at the proteome scale, including yeast two-hybrid (Y2H), affinity purification-mass spectrometry (AP-MS), protein complementation assay (PCA), and protein correlation profiling (PCP). These methods have been applied successfully to generate high-quality maps of the interactomes of humans (Hein et al., 2015; Huttlin et al., 2015; Huttlin et al., 2017; Rolland et al., 2014; Wan et al., 2015) and other metazoans (Guruharsha et al., 2011; Simonis et al., 2009).

Widely used methods for interactome mapping rely on heterologous expression or genetically manipulated cell lines, and as a consequence, existing interactome maps provide limited insight into which interactions occur in specific cell types or tissues, or under pathophysiologically relevant conditions (Snider et al., 2015). Targeted interactome mapping in tissue or cell type-specific contexts has revealed rewiring of protein interactions in select human diseases (Pankow et al., 2015; Shirasaki et al., 2012), yet fundamental questions regarding the organization of the interactome across mammalian tissues remain unanswered. Whereas large-scale efforts to profile the transcriptome, proteome, and epigenome of human tissues have been undertaken (Melé et al., 2015; Kundaje et al., 2015; Uhlén et al., 2015), a comparable resource at the interactome level is lacking.

We previously developed a high-throughput method for interactome mapping by combining size exclusion chromatography (SEC) with protein correlation profiling-stable isotope labeling by amino acids in cell culture (PCP-SILAC) (Kristensen et al., 2012). Here, we couple PCP with stable isotope labeling of mammals (SILAM) (Krüger et al., 2008; McClatchy et al., 2007) to map the interactomes of seven mouse tissues, with an accuracy comparable to the highest-quality screens in human cell lines. This resource provides the first global view of protein-protein interactions in a mammalian tissue-specific context. Our study additionally provides the first systematic interactome map in mouse, and expands the known mouse interactome by 37%, revealing over 26,000 novel interactions. The resulting *in vivo* interactome maps uncover widespread rewiring of protein interactions across physiological contexts.

## Results

### Quantitative *in vivo* interactome profiling of mouse tissues

We profiled the interactome of seven mouse tissues, including brain, heart, skeletal muscle (gastrocnemius), lung, kidney, liver, and thymus, using PCP-SILAM (Figure 1A). Organs from two male SILAM mice were pooled to minimize biological variation, and the *in vivo* interactome of each tissue was preserved by extracting and separating protein complexes under non-denaturing conditions and in the presence of excess protease and phosphorylase inhibitors. For each tissue, 55 fractions were collected, for a total of 385 unique fractions. Separate PCP fractions were pooled to generate a global reference mixture, which was spiked into all 385 fractions. Each fraction was subjected to liquid chromatography-tandem mass spectrometry (LC-MS/MS) analysis, and the resulting dataset was processed as a single experiment using MaxQuant (Cox and Mann, 2008) at a peptide and protein false discovery rate (FDR) of 1%.

**Figure 1.**
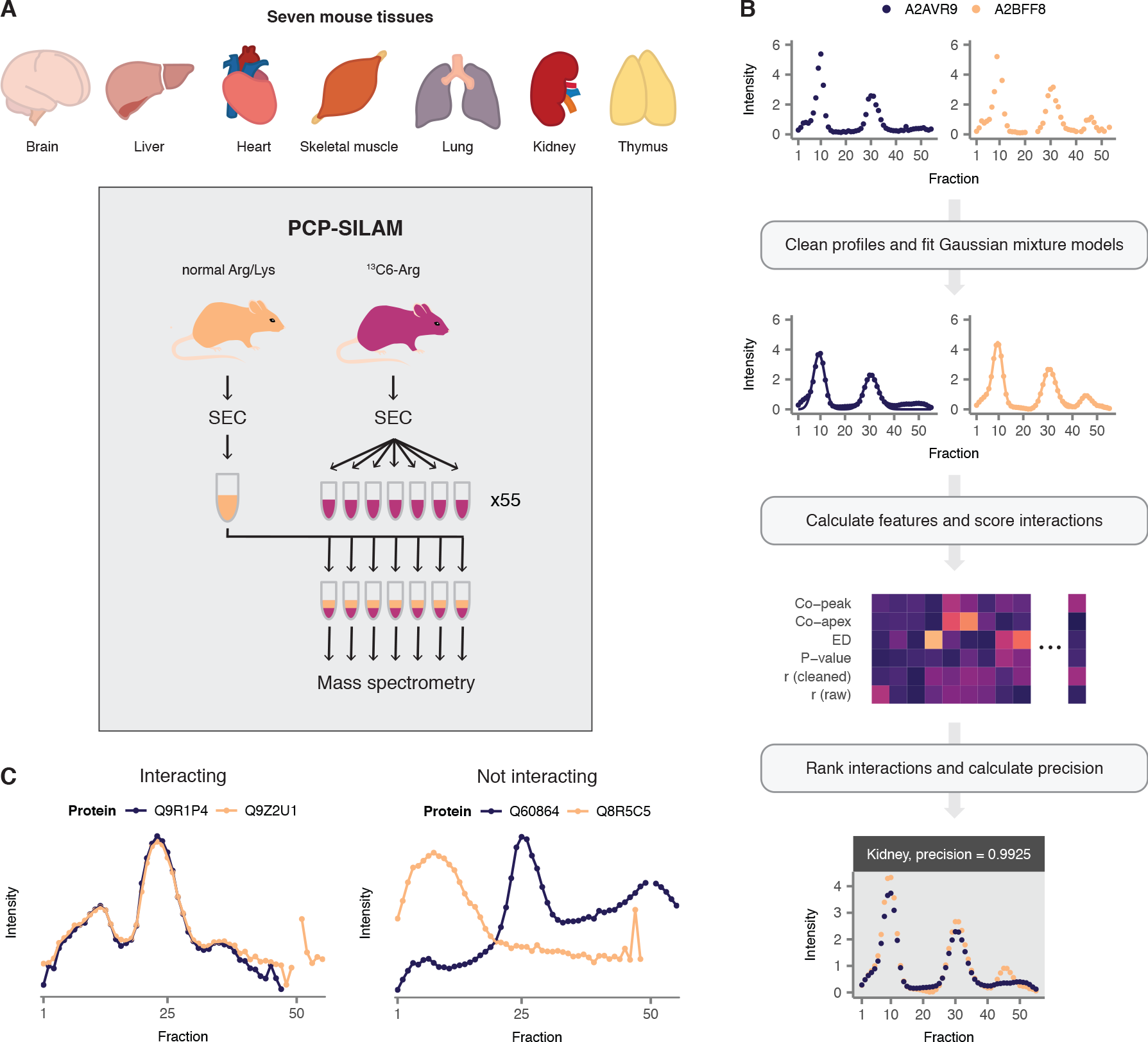
Mapping the interactomes of seven mouse tissues with PCP—SILAM. (A) PCP—SILAM workflow for interactome mapping in mouse tissues. (B) Overview of bioinformatic pipeline for recovery of interactomes from co-elution data. (C) Representative co-elution profiles for interacting and non-interacting protein pairs.

In total, 80,299 unique peptide sequences from 5,811 proteins groups were identified across all PCP fractions. Consistent with the presence of the SILAM global reference, we observed near-saturation in the quantitation of novel proteins with additional PCP fractions (Figure S1A). Our coverage of the observed tissue proteomes is comparable in depth to previous deep proteomic analyses of murine tissues (Geiger et al., 2013) (Figure S1B-C), yet contains additional functional information on interactions between proteome components. Consistent with shotgun proteomics studies (Geiger et al., 2013), the observed tissue interactomes are dominated by highly abundant proteins, which make up 75% of total protein mass (Figure S1D), while relatively few proteins are observed to be exclusively quantified within a single tissue (range 1.8%-17.4%, Figure S1E) (Kim et al., 2014). Thus, our proteomic dataset represents a rich, in-depth profile of seven murine tissue interactomes.

### Reconstruction of high-confidence mouse tissue interactomes

To derive interactomes from PCP-SILAM tissue proteome profiles, we developed PrInCE, a machine-learning pipeline for analysis of co-migration data (Figure 1B-C) (Stacey et al., 2017). PrInCE recovers PPIs using features derived exclusively from co-migration profiles, in contrast to several previous approaches, which learn jointly from co-migration data and publicly available genomic datasets such as gene coexpression profiles, phylogenetic profiles, or previously published interactions (Havugimana et al., 2012; Kastritis et al., 2017; Wan et al., 2015).

PrInCE returns a ranked list of interactions and calculates the precision at each point in the list, defined as the ratio of intra-complex interactions to intra and inter-complex interactions. To select a precision cutoff for further analysis, we compared each mouse tissue interactome to five recently published high-throughput human interactome maps (Hein et al., 2015; Huttlin et al., 2015; Huttlin et al., 2017; Rolland et al., 2014; Wan et al., 2015). At a precision of 80%, PrInCE recovered a total of 38,117 interactions across seven tissues, of which 27,004 were unique (Figure 2A). This represents 60% more unique interactions than the highest quality systematic human interactome map, at roughly equivalent precision. PrInCE additionally recovered more unique interactions than any of six literature-curated mouse interaction databases at equivalent precision (Figure S2A). The resulting interactomes, at a precision cutoff of 80%, were retained for all further analysis. Between 4,141 and 6,900 interactions were recovered for each individual tissue interactome (Figure 2B). We additionally provide the complete set of 163,665 interactions (112,539 unique) at 50% precision in Table S1.

Hierarchical clustering of the tissue interactomes, using the Jaccard index as a measure of network similarity, confirmed that these captured biologically meaningful relationships between tissues. As expected, cardiac and skeletal muscle clustered together (Figure 2C), as did thymus and lung, two tissues with a prominent immune function. Moreover, relationships between interactomes recapitulated relationships between tissues derived from the mouse transcriptome (Merkin et al., 2012) and proteome (Geiger et al., 2013) (Figure S2B). In addition, the results were similar when only interactions made by proteins quantified in all seven tissues (housekeeping proteins) were used to cluster the tissue interactomes (Figure S2C), confirming these results were not driven by tissue specificity of protein expression.

We next evaluated the biological relevance of the mouse tissue interactomes by calculating the number of interacting protein pairs in each network that share at least one annotation from the Gene Ontology (GO) (Ashburner et al., 2000). Because GO terms may be either specific or broad, enrichment relative to random pairs was calculated at three different breadths, where breadth refers to the number of proteins in the organism’s proteome annotated with that term (Simonis et al., 2009). Enrichment for functional linkage in mouse tissue interactomes was comparable to high-throughput human interactome maps at all breadths (Figure 2D and Figure S2D). Tissue interactomes also displayed comparable mRNA coexpression (Figure 2E) and enrichment for known domain-domain interactions (Figure 2F) to recent high-throughput human interactome maps, further validating the quality of the mouse tissue interactomes. In addition, we analyzed affinity purifications of the huntingtin protein from mouse brains (Shirasaki et al., 2012), and found that interacting protein pairs detected by PCP-SILAM in the mouse brain displayed highly correlated patterns of abundance across purifications (Figure S2E; *P*=2.0 × 10^−12^, Brunner-Munzel test), confirming the *in vivo* relevance of these interactions.

To further validate the ability of PCP-SILAM to identify novel interactions in mouse tissues, we focused on a putative novel interaction between the cytoskeletal protein talin and the subunits of the chaperonin containing TCP1 complex, which had not previously been reported in either mouse or human (Figure 2G). Immunoprecipitation of talin confirmed the interaction with TCP1 in mouse brain (Figure 2H). To confirm the tissue specificity of the interaction, we additionally performed immunoprecipitation in mouse liver, where talin and TCP1 complex curves displayed limited correlation, and observed limited co-purification, consistent with PCP-SILAM (Figure 2G-H). Thus, orthogonal validation confirmed the ability of PCP-SILAM to quantify dynamic rearrangements in the interactome across tissues.

**Figure 2.**
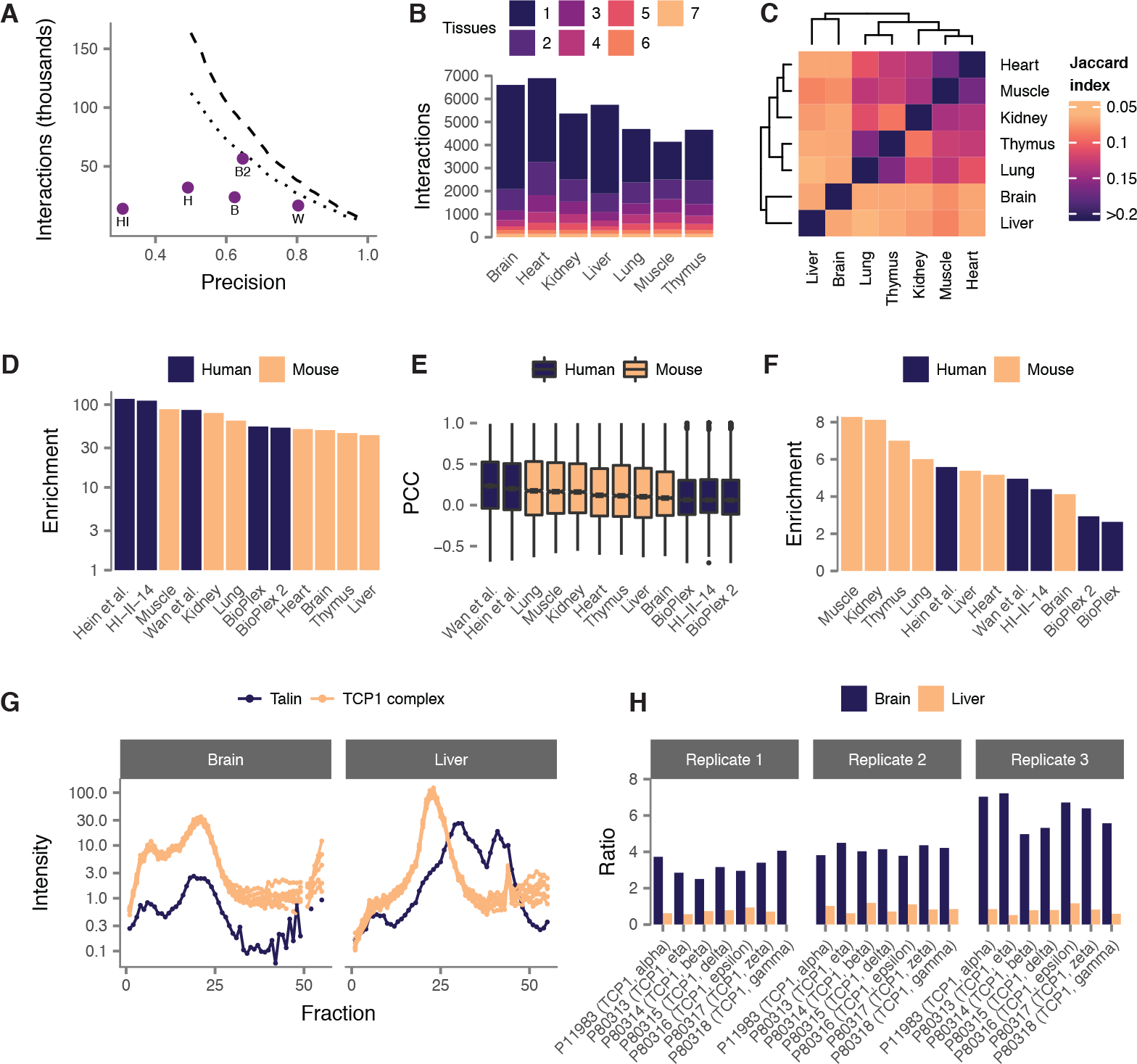
Validation of mouse tissue interactomes. (A) Precision of mouse tissue interactomes compared to five recent human high-throughput interactome maps. Dashed line shows all interactions; dotted line shows unique interactions. B, Bioplex (Huttlin et al., 2015); B2, BioPlex 2 (Huttlin et al., 2017); H, (Hein et al., 2015); HI, HI-II-14 (Rolland et al., 2014); W, (Wan et al., 2015). (B) Count and tissue specificity of interactions detected in each mouse tissue. (C) Hierarchical clustering of mouse tissue interactomes. (D) Enrichment for shared Gene Ontology terms in mouse tissue interactomes and five recent human high-throughput interactome maps. (E) Coexpression of interacting proteins in mouse tissue interactomes and human high-throughput interactomes. (F) Enrichment for domain-domain interactions in mouse tissue interactomes and human high-throughput interactomes. (G) PCP-SILAM profiles of talin and the TCP1 complex in mouse brain and liver. (H) Relative abundance of co-immunoprecipitated TCP1 complex subunits to talin in mouse brain and liver.

### Unbiased expansion of the mouse interactome by PCP-SILAM

The mouse is a ubiquitous model organism, yet its interactome has never been the subject of a systematic, proteome-scale mapping effort. We compared the PCP-SILAM mouse interactome to a literature-curated (LC) mouse interactome derived by assembling a total of 72,519 mouse PPIs that were detected in small scale experiments and reported in seven interaction databases (Table S2). Strikingly, of the 27,004 unique interactions detected by PCP-SILAM at 80% precision, only 562 (2.1%) overlap with interactions detected by small-scale experiments (Figure 3A). This overlap is significantly larger than would be expected by chance (*P* < 10^−300^, hypergeometric test), but small in magnitude, suggesting that PCP-SILAM is largely complementary to small-scale methods. Among LC PPIs, interactions detected by PCP-SILAM intersect most significantly with those detected by cosedimentation, and less with PPIs detected by two-hybrid or cross-linking approaches (Figure S3A), further supporting the notion that each assay recovers characteristic subsets of the interactome.

The remaining 26,442 unique interactions detected in this study are novel in mouse. Our proteome-scale resource therefore expands the size of the known mouse interactome by 36.5% (Figure 3B). We sought to functionally characterize the novel PPIs by comparing patterns of co-annotation between LC and PCP SILAM interactions (Figure 3C). Relative to LC interactions, mouse PPIs detected by PCP-SILAM were enriched for connections involving metabolism, translation, and protein folding. In contrast, PCP-SILAM PPIs were underrepresented in connections involving cell-cell signalling and proliferation. These findings are consistent with the expectation that PCP-SILAM would prioritize cytoplasmic over nuclear or extracellular complexes, and stable over dynamic assemblies.

We next asked whether PPIs detected by PCP-SILAM preferentially expanded existing regions of the global mouse interactome, or tended to form completely new subnetworks. To provide a global view of network topology, we applied unsupervised Markov clustering (Enright et al., 2002) to group the entire mouse interactome, including both LC and PCP-SILAM interactions, into 880 clusters of three to 4,903 proteins (Figure 3D). Of the 86 clusters containing at least one PPI detected by PCP-SILAM, 59 (68.6%) included both LC and high-throughput PPIs, while only 27 were composed exclusively of PCP-SILAM interactions. Thus, while PCP-SILAM reveals several protein communities that were altogether unknown in mouse, many interactions also expand neighborhoods of the mouse interactome previously defined by curation of small-scale experiments.

Literature-curated protein interaction datasets have been criticised on the grounds that they are biased towards a relatively small set of highly studied proteins. In agreement with a previous study in humans (Rolland et al., 2014), the mouse LC interaction dataset is dominated by interactions between well-studied proteins, at the expense of a “sparse zone” of poorly studied proteins, for which few PPIs are known (Figure 3G). High-throughput interactome mapping studies provide a means to define the architecture of the proteome independent of sociological biases, and in comparison to the LC dataset, PPIs detected by PCP-SILAM are distributed more homogeneously with respect to the number of publications (Figure 3G). However, we anticipated that the untargeted nature of our *in vivo* approach could result in a reduced capacity to detect interactions for lowly expressed proteins, relative to targeted approaches. Consistent with this expectation, PPIs detected by PCP-SILAM display a moderate bias towards more abundant proteins, whereas interactions detected by small-scale experiments largely do not (Figure S3C).

To date, large-scale interactome mapping projects have typically been performed in yeast cells or cell lines (Snider et al., 2015). However, many biologically relevant PPIs may not occur within these systems. We therefore hypothesized that mapping the *in vivo* interactome in physiological contexts, such as in individual tissues, could preferentially reveal novel, context-specific interactions. Consistent with this hypothesis, we found that tissue-specific interactions were significantly less likely to have been catalogued in literature-curated interaction databases (Figure S3B; *P* = 5.8 × 10^−145^, Kendall rank correlation). We also investigated whether mapping interactomes in mouse tissues could preferentially provide insights into interactions involving proteins whose functions are poorly understood. PCP-SILAM identified interacting partners for 557 proteins of unknown function (Table S3A), and these interactions were significantly more tissue-specific than the interactome average (Figure 3E; *P*=4.4 × 10^−22^, Kendall rank correlation). Similarly, we mapped interactions involving 666 proteins for which no interacting partners had previously been detected (interactome “orphans,” Table S3B) (Kotlyar et al., 2015), and found that these interactions likewise displayed a significant trend towards increasing tissue specificity (Figure 3F; *P*=2.9 × 10^−54^, Kendall rank correlation). Thus, mapping the *in vivo* interactome of mammalian tissues can help shed light on poorly studied components of the proteome.

**Figure 3.**
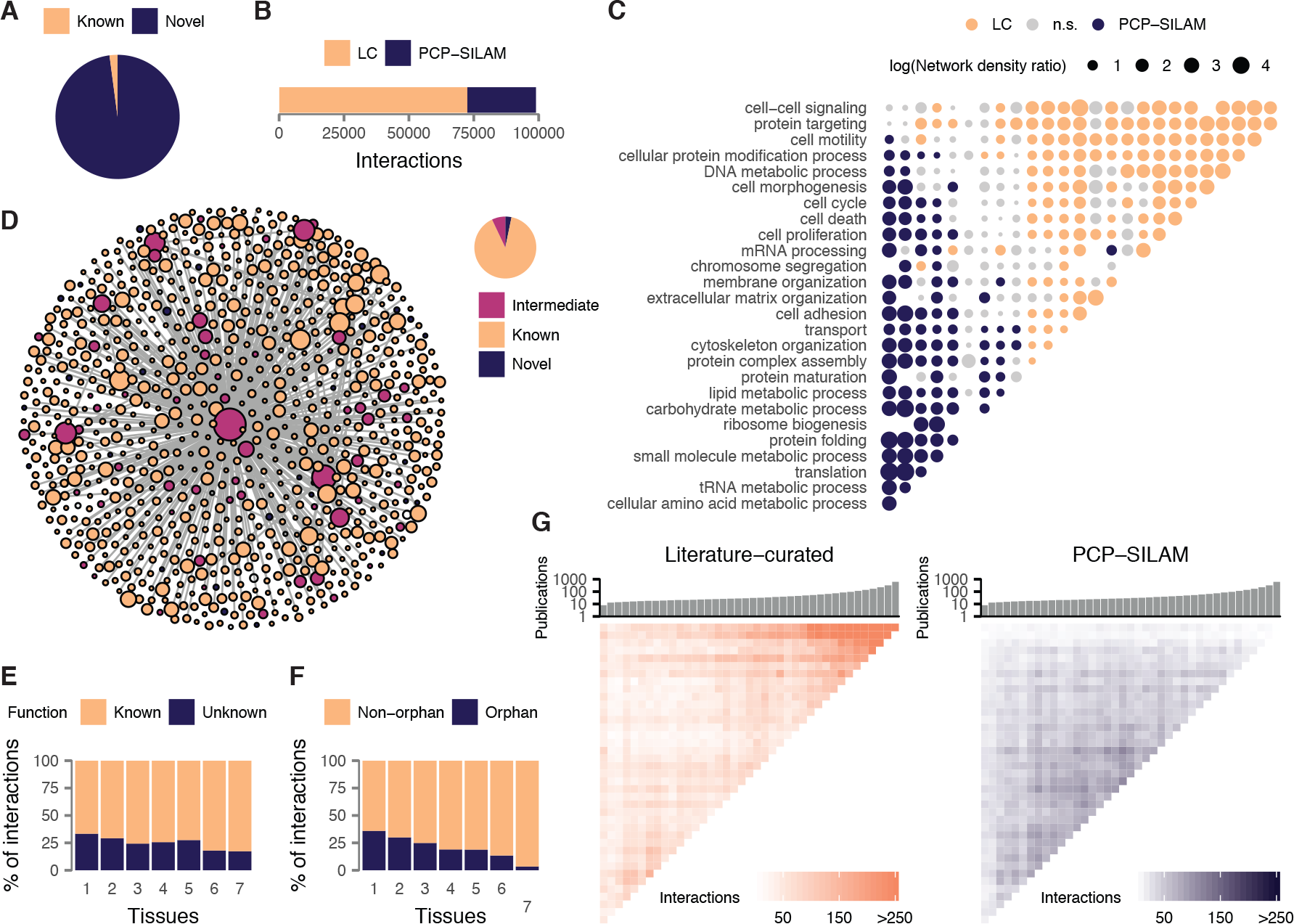
Unbiased expansion of the literature-curated mouse interactome by PCP-SILAM. (A) Proportion of previously known (2.1%) and novel (97.9%) unique interactions observed in PCP-SILAM mouse tissue interactomes. (B) Size of the known mouse interactome before and after the present study. (C) Comparison of interacting protein co-annotation within and across biological processes between PCP-SILAM and LC interactions. (D) Markov clustering of the global mouse interactome, and proportion of protein communities containing exclusively known interactions, exclusively novel interactions, or both. (E) Proportion of interactions involving proteins of unknown function for interactions detected in one to seven tissues. (F) Proportion of interactions involving proteins for which no interactions were previously known (interactome “orphans”) for interactions detected in one to seven tissues. (G) Number of interactions between proteins binned by number of publications, and ordered along both axes. Histogram shows the median number of publications in each bin.

### Widespread interactome rewiring limits the accuracy of tissue-specific interactome prediction

In the absence of experimental tissue or cell type-specific interactomes, computational methods have been developed to reconstruct context-specific molecular interaction networks, with a view to understanding network perturbations in tissue-specific pathologies (Greene et al., 2015; Marbach et al., 2016). The most widely used strategy for context-specific interactome prediction proceeds from the notion that the protein products of two genes can only interact in a given context if these genes are both expressed. Thus, gene or protein expression data is overlaid onto a static interactome, and the subset of the network whose nodes are expressed above a certain threshold is extracted to generate the context-specific interactome (Figure 4A) (Bossi and Lehner, 2009; Buljan et al., 2012; de Lichtenberg et al., 2005). A second strategy is to construct tissue-specific gene coexpression networks, which suggest functional association in a given tissue, if not necessarily physical interaction (Parikshak et al., 2013; Voineagu et al., 2011).

Using our PCP-SILAM mouse tissue interactomes as a reference, we investigated the accuracy of these approaches in predicting tissue-specific interactomes. Surprisingly, tissue interactomes predicted based on gene expression were only two to four-fold enriched for experimentally detected interactions, relative to randomized networks (range 2.5-4.1, Figure 4B). This overlap is highly significant (all *P* ≤9.9 × 10^−47^, hypergeometric test), but remarkably small in magnitude. Similar results were observed for tissue-specific coexpression networks (Figure 4B), consistent with previous findings that gene coexpression is a relatively poor predictor of physical interaction (Kühner et al., 2009). Thus, neither tissue-specific gene expression nor coexpression are sufficient to accurately predict tissue-specific physical PPIs.

Predicted tissue interactomes also differed markedly in their topology from experimentally determined interactomes. In interactome networks, the most highly connected (“hub”) proteins are slow-evolving and physiologically indispensable (Fraser et al., 2002; Jeong et al., 2001). The hub proteins of predicted tissue interactomes were highly consistent across tissues, with 61% to 79% of hubs in each tissue common to all five predicted networks (Figure S4A). However, we found that hubs are much less ubiquitous, with only two hubs shared across all five tissues (Figure S4A). Moreover, the identities of the hub proteins themselves in each tissue were also poorly predicted, with an overlap of only 9-17% between hub proteins in predicted and *in vivo* interactomes (Figure 4C). More generally, the number of interactions a protein makes (its degree) was much more variable between tissues than in predicted networks (Figure S4B). Similarly, PCP-SILAM revealed that proteins display a much greater tendency to interact with different partners across tissues than was predicted by gene expression alone (Figure 4D and S4C).

Taken together, these analyses reveal widespread rewiring of interactome networks across mouse tissues beyond what is apparent from gene expression alone, affecting both the specific interactors of individual proteins and the global topological properties of physiological interactomes. Critically, the observation that a pair of proteins can interact in at least one context does not imply that their expression in a second context is a sufficient condition to reproduce the interaction. For example, PCP-SILAM correctly identified the known interaction between the F-actin-capping protein (CapZ) and the CapZ-interacting protein (CapZIP or Rcsd1) (Edwards et al., 2014; Eyers et al., 2005). However, the interaction was specific to heart (98.8% precision) and skeletal muscle (78.8% precision), despite robust expression of the interacting proteins in all seven tissues (Figure 4F). This pattern of tissue specificity is consistent with the proposed function of the interaction in muscle contraction (Eyers et al., 2005), but could not have been predicted on the basis of protein abundance alone. To evaluate whether predicted interactomes are depleted for tissue-specific interactions more systematically, we randomized interaction networks for each tissue separately, and calculated the total number of interactions found across one to seven randomized tissue interactomes. Consistent with the expectation that gene expression alone would underestimate the degree of interactome variability across tissues, predicted interactomes were significantly depleted for the most tissue-specific interactions (i.e., those found in only a single tissue), relative to PCP-SILAM interactomes (Figure 4E and S4D; odds ratio = 0.14, *P* = 9.7 × 10^−10^, Z test). In summary, gene expression alone is insufficient to explain the observed rewiring of protein interactions across tissues, limiting the acuracy of existing methods to predict context-specific interactomes.

**Figure 4.**
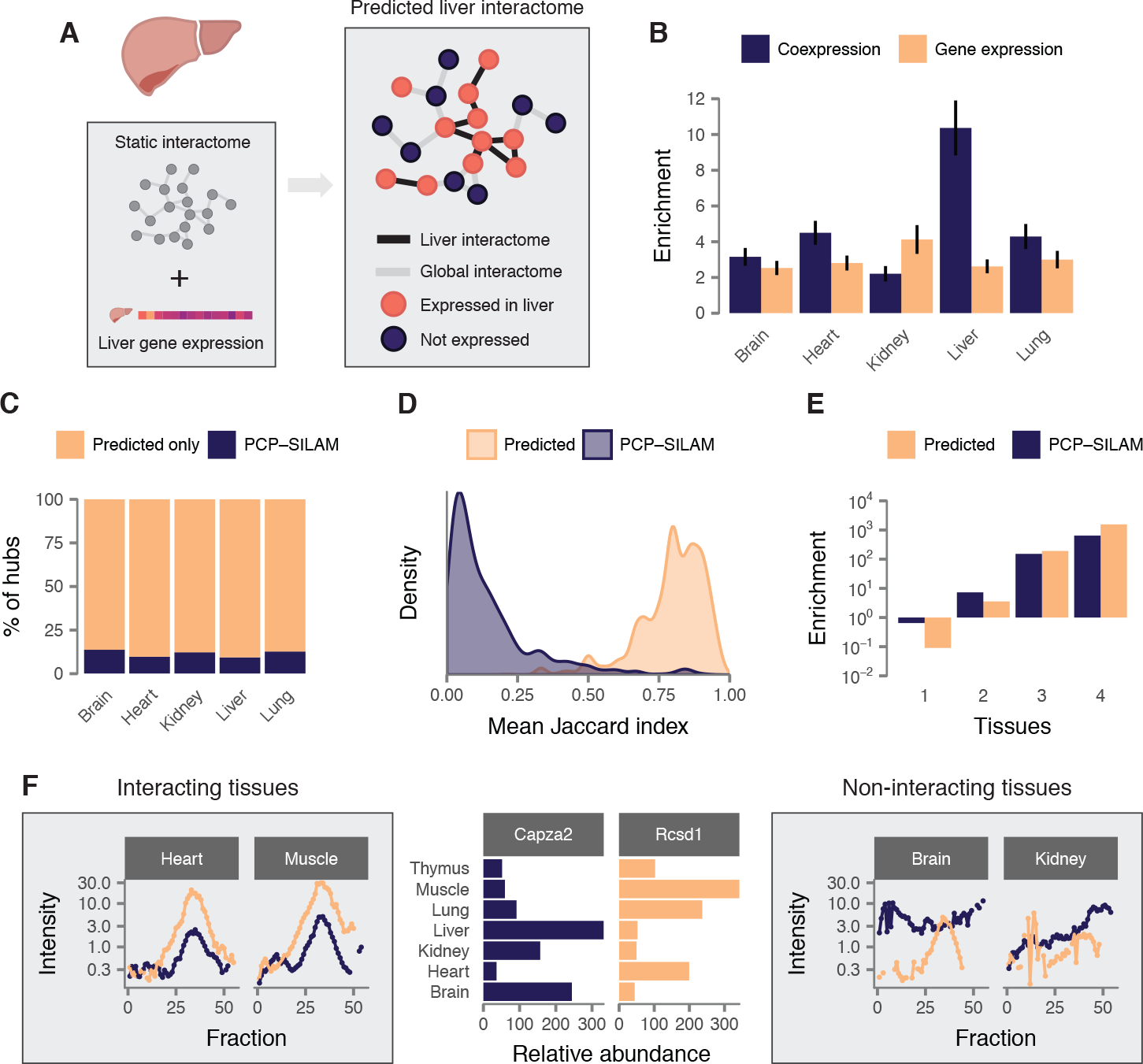
Widespread interactome rewiring limits accuracy of tissue-specific interactome prediction. (A) Overview of tissue-specific interactome prediction by integration of static interactome maps with tissue-specific gene expression profiles. (B) Overlap between PCP-SILAM mouse tissue interactomes and tissue-specific gene coexpression networks or tissue interactomes predicted based on gene expression, relative to rewired networks. (C) Most PCP-SILAM tissue interactome hubs are not hubs in the corresponding predicted tissue interactomes. (D) Protein rewiring across tissue interactomes in predicted and PCP-SILAM tissue interactomes, quantified by the mean Jaccard index across all pairs of tissues. (E) Enrichment for interactions found in one to four tissues, relative to randomized tissue interactomes. PCP-SILAM profiles reveal Rcsd1 and Capza2 interact in heart and muscle, but not brain or kidney, despite expression in all seven tissues. Left, PCP-SILAM chromatograms for Rcsd1 (orange) and Capza2 (purple) in heart and muscle. Middle, summed SILAM ratios for Capza2 and Rcsd1 in each tissue. Right, PCP-SILAM chromatograms in brain and kidney.

### Evolution of interactions in mammalian tissues

Extrapolation of physical interactions detected in one organism to orthologous pairs of proteins in a different organism has been widely used to predict the interactomes of non-model organisms, or increase coverage of the human interactome (Li et al., 2017; Matthews et al., 2001; Wan et al., 2015; Yu et al., 2004). However, many tissues execute specialized biological functions that are poorly conserved between organisms. We hypothesized that more tissue-specific interactions would show less evidence of evolutionary conservation than interactions that occur across many tissues. Examining literature-curated interactions for three model organisms, and five recent human high-throughput interactome maps, we found that tissue-specific interactions were less likely to be evolutionarily conserved (Figures 5A and S4A; *P* = 6.9 × 10^−152^ and *P* <10^−300^ for model organisms and humans respectively, Fisher integration of Kendall rank correlation). Protein pairs that made tissue-specific interactions likewise showed less similar evolutionary rates (Figure 5B; *P* = 5.1 × 10^−18^, Kendall rank correlation), and had more distinct phylogenetic profiles (Figure 5C; *P* = 9.5 × 10^−55^), relative to universal interactions. Thus, both experimental and genomic data highlight the evolutionary novelty of tissue-specific interactions.

We next asked whether tissue-specific interactions predominantly arise from tissue-specific rewiring of ancient proteins, or whether they instead involve evolutionarily young proteins. Relative to universal interactions, tissue-specific interactions disproportionately involved younger proteins (Figure S5B; *P* = 8.3 × 10^−81^). Furthermore, ancient proteins had more conserved interaction partners across tissues, whereas younger proteins were disproportionately rewired between tissue interactomes (Figure S5C; *P* =2.6 × 10^−10^), suggesting that rewiring of ancient proteins is insufficient to explain the evolution of novel, tissue-specific interactions.

Previous analyses of predicted tissue interactomes identified extensive interactions between proteins expressed in only a subset of tissues and those expressed in all tissues (housekeeping proteins), proposing a model wherein tissue-specific functions arise by recruiting core cellular processes (Bossi and Lehner, 2009). Motivated by our observation that interactome rewiring is poorly predicted by tissue-specific gene expression, we investigated whether PCP-SILAM data supported a model of extensive cross-talk between housekeeping and tissue-specific proteins, or one in which tissue-specific functions are mediated independently of core cellular processes. To quantify the extent of cross-talk between housekeeping and tissue-specific proteins, we randomized each tissue interactome, using a degree-preserving method to control for network topology (Maslov and Sneppen, 2002), and compared the number of observed interactions between proteins at each level of tissue specificity to random expectation. In every tissue, housekeeping proteins displayed highly significant enrichment for interactions with other housekeeping proteins, offering robust statistical support for the existence of core cellular modules within the interactome (Figure 5D). In contrast, we observed systematic depletion of interactions between tissue-specific and housekeeping proteins, particularly when aggregating results across all seven tissues (Figure 5E). Experimental tissue interactome mapping therefore reveals that evolutionarily novel tissue-specific interactions accomplish tissue-specific functions largely independent of the core modules of universal interactions.

Taken together, these analyses contrast two systems: core cellular modules present across all mouse tissues, and accessory modules that execute specialized functions within individual tissues. The former involves ancient proteins that have co-evolved over long evolutionary timeframes, and whose interaction partners are preserved across species and tissues. In contrast, tissue-specific interactions are less often conserved in other species and disproportionately involve younger proteins. Remarkably, we observe suppression of cross-talk between these two systems, with significant depletion of interactions between tissue-specific and universal proteins. Importantly, the opposite conclusion is reached when relying solely on tissue interactomes predicted based on gene expression, highlighting the importance of experimental interactome mapping to develop accurate systems-level models of biological processes.

**Figure 5.**
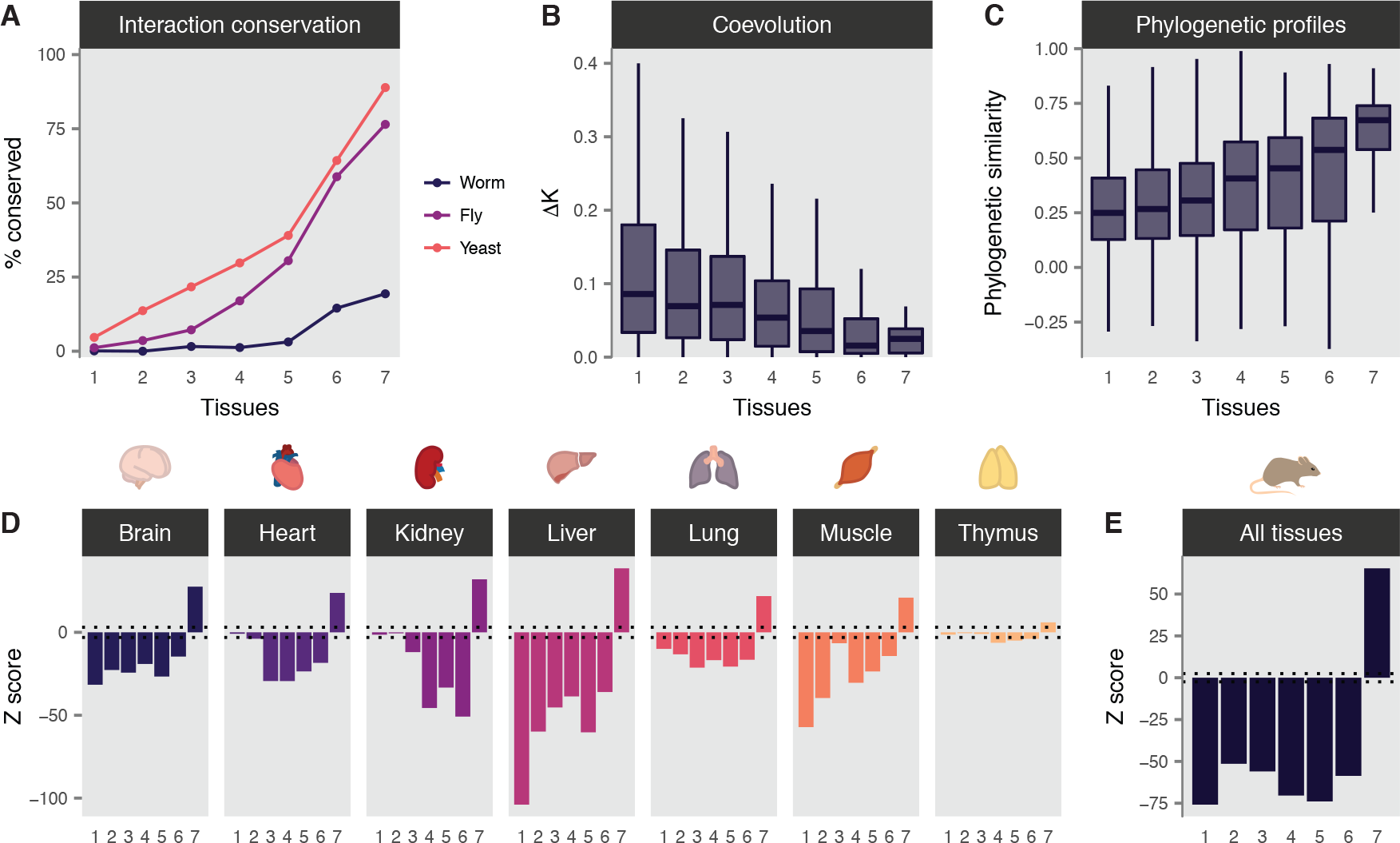
Evolution of mammalian tissue interactomes. (A) Proportion of mouse interactions conserved in worm, fly, and yeast for interactions detected in one to seven tissues. (B) Difference in evolutionary rates between mouse interactions detected in one to seven tissues. (C) Correlation in phylogenetic profiles between mouse interactions detected in one to seven tissues. (D-E) Z scores for the statistical significance of interactions between housekeeping proteins and proteins quantified in one to seven tissues, relative to randomized networks in each mouse tissue interactome (D) and aggregated across mouse tissues (E).

### Tight regulation of tissue-specific interaction rewiring

Having established that evolutionarily ancient proteins have significantly more preserved interaction partners across tissues than the interactome average, we sought to further characterize the properties of proteins whose interactions are disproportionately rewired in a tissue-specific manner. We quantified the degree to which a protein was rewired across tissues by calculating the mean Jaccard index of its interacting partners across all pairs of tissues (Figure S6A). Proteins with a higher Jaccard index participate in interactions that are preserved across mouse tissues, whereas proteins with a low Jaccard index have interaction partners that are more rewired.

As expected, members of known protein complexes were significantly less rewired across tissues (i.e., they had a greater Jaccard index; Figure 6A, *P* = 5.9 × 10^−7^, one-tailed Brunner-Munzel test). Surprisingly, rewired proteins were not enriched for any functional annotation. However, proteins without any known function at all were significantly more rewired (Figure 6B, *P* = 1.0 × 10^−3^). Examining protein structural features, we found that proteins containing a globular domain were significantly less rewired than the interactome average (Figure 6C, *P* = 1.2 × 10^−3^). In contrast, intrinsically disordered proteins were significantly more rewired (Figure 6D, *P* = 6.4 × 10^−4^). Disordered segments of proteins often contain short peptide interaction motifs that can be bound by globular domains (Davey et al., 2012), and proteins containing such linear motifs were likewise significantly more rewired (Figure 6E, *P* = 5.6 × 10^−3^). Together, these findings suggest that binding motifs embedded within intrinsically disordered regions facilitate the rewiring of protein interaction partners across mammalian tissues.

**Figure 6.**
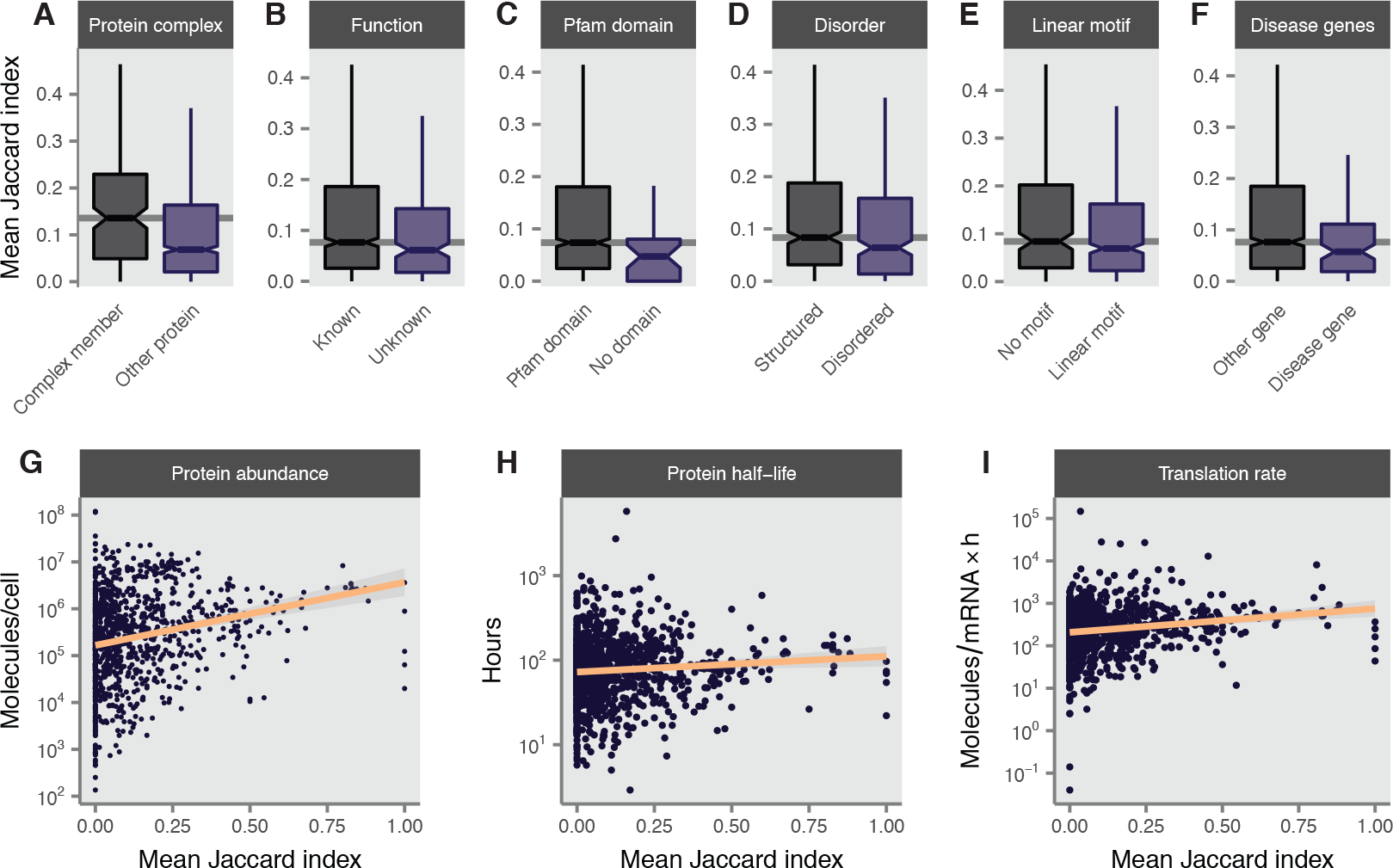
Tight regulation of interaction rewiring. (A) Members of known protein complexes display a lesser degree of interaction rewiring across tissue interactomes. (B) Proteins of unknown function display a greater degree of interaction rewiring across tissue interactomes. (C) Proteins without a globular protein domain display a greater degree of interaction rewiring across tissue interac tomes. (D) Intrinsically disordered proteins display a greater degree of interaction rewiring across tissue interactomes. (E) Proteins containing linear protein-binding motifs display a greater degree of interaction rewiring across tissue interactomes. (F) Mouse disease genes display a greater degree of interaction rewiring across tissue interactomes. (G-I) Rewired proteins are characterized by low abundance (G), short half-lives (H), and slow translation rates (I).

Disordered proteins are frequently involved in signaling pathways or mediate regulatory functions (Ward et al., 2004). We therefore hypothesized that rewiring of PPIs across tissues could facilitate tissue-specific signalling processes. Consistent with this hypothesis, tissue-specific interactions were significantly more likely to involve protein kinases, transcription factors, and cell surface protein receptors (Figure S6B-D, *P* = 6.0 × 10^−6^, 4.6 × 10^−5^, and 1.7 × 10 ^−6^ respectively, Kendall rank correlation). In addition, we calculated the betweenness centrality of each interaction, defined as the number of shortest paths across the network that pass through each edge. Edges with a high betweenness centrality in interactome networks are associated with information flow across the network (Yan et al., 2016), and in agreement with this notion, we found tissue-specific interactions to have a significantly higher centrality than universal interactions (Figure S6E, *P* < 10^−300^, Kendall rank correlation). Thus, both molecular and network topological perspectives highlight the key role of tissue-specific interactions in propagating biological information within tissue-specific pathways.

Within the cell, precise coordination of macromolecular interactions is required for accurate transmission of biological information. We therefore hypothesized that proteins whose interaction partners are highly variable across physiological contexts would be subject to tight regulatory mechanisms, and asked whether specific cellular strategies regulate the availability of rewired proteins. mRNAs encoding rewired proteins were expressed at lower levels, had shorter half-lives, and were transcribed at slower rates (Figure S6F-H; *P* = 7.4 × 10^−12^, 3.6 × 10^−4^, and 4.3 × 10^−12^ respectively, Spearman rank correlation) than proteins with stable interaction partners. Rewired proteins themselves were also less abundant (Figure 6G, *P* = 1.6 × 10^−20^), and this difference in abundance was controlled both by a reduced rate of translation and increased rate of degradation (Figure 6H-I, *P* = 4.8 × 10^−8^ and 1.0 × 10^−4^, respectively), suggesting multiple cellular mechanisms converge to tightly regulate the abundance of proteins whose interacting partners are rewired across tissues. Disordered proteins are known to be tightly regulated (Gsponer et al., 2008), but partial correlation analysis confirmed the tight regulation of rewired proteins was independent of protein disorder for all outcomes except protein half-life (all *P* ≤ 3.9 × 10^−3^, partial Spearman correlation). Given this tight regulation of rewired proteins, we further asked whether proteins whose interaction partners are highly rewired between tissues were associated with deleterious phenotypes. Remarkably, we found that mouse disease genes were significantly more rewired across tissues than the interactome average (Figure 6F, *P* = 8.2 × 10^−3^, one-tailed Brunner-Munzel test).

Taken together, these analyses highlight the role of protein-binding motifs within disordered regions in mediating interaction rewiring across physiological contexts. The resulting tissue-specific interactions are associated with transmission of biological information in tissue-specific signalling pathways. Highly rewired proteins are subject to tight regulation by multiple convergent cellular mechanisms, perhaps to ensure the fidelity of biological information flow, and the elevated rate of interaction rewiring among disease genes implicates dysfunction of this regulatory cascade in disease pathophysiology.

## Discussion

Defining the protein-protein interactome is essential to revealing the molecular origins of cellular processes. However, earlier efforts produced static interactome maps that are fundamentally limited with respect to understanding interaction dynamics across tissue or cell type-specific contexts. By applying PCP-SILAM to map the *in vivo* interactome of seven mouse tissues, we provide a systematic, proteome-scale resource to understand the dynamic physiological interactome. Multiple functional genomics measures, including functional co-annotation, mRNA co-expression, and domain-domain interactions, indicate that our experimental and bioinformatic pipeline mapped tissue-specific interaction networks with an acuracy comparable to the highest-quality human high-throughput screens (Hein et al., 2015; Huttlin et al., 2015; Huttlin et al., 2017; Rolland et al., 2014; Wan et al., 2015).

In the absence of high-throughput methods to define context-specific *in vivo* interactomes, computational methods have been developed to predict interactome rewiring based on gene expression (Bossi and Lehner, 2009; Buljan et al., 2012; de Lichtenberg et al., 2005). We find that widespread interactome rewiring limits the accuracy of tissue interactome predictions based on tissue-specific patterns of gene expression or coexpression. The effect of this rewiring extends beyond the mere identities of the interactors in each tissue to the global topological properties of tissue-specific interactomes. This finding reinforces conclusions from targeted studies, which have revealed marked dissimilarities in PPIs across cell lines (Floyd et al., 2016; Jäger et al., 2011), between cellular compartments (Markmiller et al., 2018), in response to cellular stimulation (Kristensen et al., 2012; Scott et al., 2015), or in disease-relevant contexts (Pankow et al., 2015; Shirasaki et al., 2012). In yeast, widespread interaction rewiring has been observed in response to environmental perturbagens (Celaj et al., 2017). Our systematic screen builds on these findings, revealing interactome rewiring at a much larger scale and across healthy mammalian tissues. As an example of an insight into interactome organization that is not apparent from gene expression-guided predictions of context-specific interactomes, we identify systematic suppression of cross-talk between the core housekeeping interactome and tissue-specific modules, whereas the opposite conclusion had been reached in analyses of predicted tissue interactomes (Bossi and Lehner, 2009).

Evolutionary analyses of mouse tissue interactomes contrast core cellular modules composed of evolutionarily ancient, housekeeping proteins that are connected via universal interactions with evolutionarily recent proteins that interact in a more tissue-specific manner and are associated with cellular signalling. Intrinsically disordered proteins are particularly predisposed to interaction rewiring across tissues, consistent with the notion that these proteins can adopt new interacting partners over relatively rapid evolutionary timescales (Hultqvist et al., 2017). Our findings linking proteins containing linear motifs or intrinsically disordered regions to an *in vivo* program of interactome rewiring substantiate previous bioinformatic or *in vitro* analyses suggesting that alternative splicing of disordered protein-coding exons can facilitate interactome remodeling (Buljan et al., 2012; Ellis et al., 2012; Romero et al., 2006). Proteins whose interactions are highly rewired across tissues are subject to tight cellular regulation, and are implicated in disease, suggesting dysfunction of this regulatory program is at the heart of many deleterious phenotypes.

The quantitative proteomic method presented in this study, PCP-SILAM, has the advantage of being an untargeted and relatively unbiased technique, apart from its moderate bias towards proteins of higher cellular abundance. As a high-throughput technique for *in vivo* interactome mapping, PCP-SILAM is uniquely suited to the simultaneous discovery of novel interactions and quantification of their dynamics across tissues. Interrogation of *in vivo* interactomes with PCP-SILAM maps new regions and functional classes within the mouse interactome, and places poorly-studied mouse proteins into tissue-specific functional contexts, suggesting PCP-SILAM will be a valuable method to shed light on poorly understood components of the proteome.

The biological picture that emerges from our systematic map of seven mouse tissue interactomes is one of widespread interactome rewiring, and this rewiring may be a crucial mediator of cellular and organismal phenotype. The extent of interactome rewiring observed across healthy mammalian tissues in this study indicates that a complete understanding of the human interactome will require experimental definition of the context-specific interactome networks across cell types and tissues, and their dynamic changes in response to cellular stimulation, differentiation, and disease. Our study serves as a first step towards this goal and provides a foundation for building a systems-level understanding of the mechanistic roles interactome rewiring plays in health and disease.

## Acknowledgements

M.A.S. is supported by a CIHR Vanier Canada Graduate Scholarship, an Izaak Walton Killam Memorial Pre-Doctoral Fellowship, a UBC Four Year Fellowship, and a Vancouver Coastal Health-CIHR-UBC MD/PhD Studentship. N.E.S. was supported by a Michael Smith Foundation Post-doctoral Fellowship (award #5363) and a National Health and Medical Research Council of Australia (NHMRC) Overseas (Biomedical) Fellow (APP1037373). This work was supported by funding from Genome Canada and Genome British Columbia (project 214PRO) and the Canadian Institutes of Health Research (MOP77688) to L.J.F. The authors thank Alison McAfee for constructive reading of the manuscript.

## Methods

### Generation of SILAM tissues

Protocols for the generation of SILAM labelled and unlabelled tissues were approved by the University of British Columbia Animal Care Committee in accordance with international guidelines (protocol number: A13-0094). SILAM colonies and unlabelled littermate controls were generated according to the approach of Krüger et al. (Krüger et al., 2008; Zanivan et al., 2012). Briefly, two sets of female and male littermate C57BL/6 mice (Charles River Laboratories) aged 8 weeks were segregated into separate cages and female mice feed either a SILAC chow diet (8 g ^13^C6-Lysine/kg, Cambridge Isotope Laboratories, Andover, MA) or unlabelled matched chow diet differing only in the incorporation of ^13^C6-Lysine. After 10 weeks of SILAC feeding, male mice were introduced to F0 SILAM female to allow mating. The resulting F1 SILAM mice were allowed to develop to 8 weeks and subjected to another round of mating to generate F2 SILAM mice. F2 SILAM mice were allowed to develop to 8 weeks prior to tissue isolation and protein complex extraction. In parallel to the generation of SILAM mice, unlabelled mice were derived from littermates of the F0 SILAM breed and allowed to develop to 8 weeks.

### SILAM incorporation monitoring

SILAM incorporation rates were monitored in female mice during the generation of F1 and F2 litters. For blood sample collection, mice were anesthetized with isoflurane and ~20 *μ*L of blood was collected by tail snipping every month during SILAM feeding. To measure isotope incorporation, the medulla oblongata was collected during animal termination of F0, F1 and F2 mice. Blood and medulla oblongata samples for SILAM incorporation analysis were snap frozen in liquid nitrogen and stored at −80°C. Samples were boiled in 1% sodium deoxycholate, digested, and quantified as described below.

### Tissue harvesting

For each tissue, the organs from two 8-week old C57BL/6 Male mice were harvested and pooled for downstream processing. C57BL/6 mice were terminally anesthetized with isoflurane and after the loss of corneal reflexes moved to a chilled surgical platform. To limit blood contamination within tissues and inhibit phosphatase and protease activity, the heart was exposed by Y-incision and mice perfused with 50 mL of ice-cold size exclusion chromatography (SEC) mobile phase [50 mM KCl, 50 mM NaCH_3_COO, pH 7.2, containing 2 X Complete protease inhibitor cocktail without EDTA (Roche) and 2 X Halt^TM^ Protease and Phosphatase Inhibitor Cocktail (Thermo Scientific, San Jose CA)]. Perfusate was introduced into the left ventricle by précising the ventricle wall with a needle while the right ventricle was cut to allow drainage. Upon complete blanching of the liver, the seven tissues of interest (heart, brain, thymus, liver, kidney, skeletal muscle, and lung) were removed, rinsed with ice-cold SEC buffer and placed into ice-cold SEC containing 2 X Complete protease inhibitor cocktail without EDTA and 2 X Halt^TM^ Protease and Phosphatase Inhibitor Cocktail. Tissues were further cut into smaller pieces to enhance accessibility of inhibitors and placed on ice.

### Preparation of cytoplasmic complexes for PCP separation

Complex preparation of dissected tissues and size exclusion chromatography were performed as described previously (Kristensen et al., 2012), with minor modifications. Briefly, tissues samples were lysed using a Dounce homogenizer with 200 strokes of a loose pestle followed by 200 strokes with a tight pestle. Lysates were ultracentrifuged at 100,000 relative centrifugal force (r.c.f.) for 15 min at 4°C to remove insoluble material and to partially deplete highly abundant ribosomes. Large molecular weight complexes were then concentrated using 100,000 Da molecular weight cut-off spin columns (Sartorius Stedim, Goettingen, Germany). Five milligrams of total filtered protein was then immediately loaded onto a chromatography system consisting of two 300 × 7.8 mm BioSep-4000 Columns (Phenomenex, Torrance, CA) equilibrated with SEC mobile phase and separated into 80 fractions by a 1200 Series semi-preparative HPLC (Agilent Technologies, Santa Clara, CA) at a flow rate of 0.5 mL/min at 8°C. The HPLC consisted of a G1310A isocratic pump, G7725i manual injector, and G1364 fraction collector with a G1330 thermostat. Fractions 1 to 55 corresponded to molecular weights from 2 MDa to 100 KDa, as determined by the use of common SEC standards thyroglobulin, apoferritin and bovine serum albumin (Sigma Aldrich). Each tissue was separated independently by SEC for both labelled and unlabelled samples. Fractions 1 to 55 from the seven heavy-labelled tissue preparations were pooled and served as an internal reference allowing the comparison between and across all samples. The pooled reference was spiked into each of the corresponding light fractions at a volume of 1:0.75 (light to heavy).

### In-solution digestion of PCP-SEC samples

Individual PCP-SILAM samples were prepared using in-solution digestion as previously described (Rogers et al., 2010). Briefly, sodium deoxycholate was added to each fraction to a final concentration of 1.0% (w/v) and samples boiled for 5 min. Boiled samples were allowed to cool to RT then reduced for 1 hour with 10 mM dithiothreitol (DTT) at room temperature. Samples were then alkylated for 1 h with 20 mM iodoacetamide (IAA) in the dark at room temperature and excess IAA quenched with 40mM DTT for 20 min. Two micrograms of Lys-C (Wako) was added to each fraction and samples were incubated overnight at 37° C with shaking. Samples were acidified to pH < 3 with acetic acid to precipitate deoxycholic acid, which was then removed by centrifugation at 16,000 r.c.f. for 20 min. To ensure the removal of particulate matter, peptide digests were further clarified using Unifilter 800 Whatman filter plates (GE Healthcare Life Sciences). The resulting peptide supernatant was purified using self-made Stop-and-go-extraction tips (StageTips) (Rappsilber et al., 2007) composed of C18 Empore material (3M) packed in to 200 *μ*L pipette tips. Prior to addition of the peptide solution, StageTips were conditioned with methanol, followed by 80% MeCN, 0.1% formic acid (Buffer B), then 0.1% formic acid (Buffer A). Peptide supernatants were loaded onto columns and washed with three bed volumes of Buffer A. Peptide samples were stored directly on stage tip at 4°C until required, when they were eluted with (Buffer B) directly into a HPLC autosampler plate and dried using a vacuum concentrator.

### Liquid chromatography and mass spectrometry analysis

Prior to LC—MS/MS analysis, samples were resuspended in 15 *μ*L Buffer A. LC—MS/MS was performed on an EASY-nLC1000 system (Thermo Scientific) coupled to a Q-Exactive mass spectrometer (Thermo Scientific) for biological replicates of the quantitative studies. LC—MS/MS was accomplished using a two-column system in which samples were concentrated prior to separation on a 2-cm-long, 100-im-inner diameter fused silica trap column containing 5.0-*μ*m Aqua C18 beads (Phenomenex) and then separated using an in-house packed C18 analytical 75 *μ*m inner diameter, 360 *μ*m outer diameter column composed of 35 cm ReproSil-Pur C18 AQ 1.9 *μ*m (Dr. Maisch, Ammerbuch-Entringen, Germany) column. Samples were concentrated onto the trap for 5 min using 100% Buffer A at 5 *μ*L/min after which the gradient was altered from 100% Buffer A to 40% Buffer B over 180 min at 250 nL/min with the eluting peptides infused directly into the mass spectrometers via nESI. The Q-Exactive was operated in a data-dependent manner using Xcalibur v2.2 (Thermo Scientific) with the top ten most intense multiply-charged ions above a 5% underfill ratio from MS1 scans (resolution 70,000; 350-2,000 m/z, AGC target of 3 × 10^6^) selected for HCD MS-MS events (resolution 17.5k AGC target of 1 × 10^6^ with a maximum injection time of 60 ms, NCE 28 with 20% stepping) with 25 s dynamic exclusion enabled.

### Proteomic data analysis

MaxQuant (version 1.5.5.1) (Cox and Mann, 2008) was used for identification and quantification of the resulting experiments, with the resulting cytoplasmic and membrane biological replicates searched together to ensure an accurate global false discovery rate in accordance with the work of Schaab et al. (Schaab et al., 2012). Database searching was carried out against the UniProt Mus musculus database (downloaded August 17, 2014; 49,235 entries) with the following search parameters: carbamidomethylation of cysteine as a fixed modification, oxidation of methionine, acetylation of protein N-termini, semi-LysC cleavage with a maximum of two missed cleavages. A multiplicity of two was used, denoting the SILAM amino acid combinations (light lysine and heavy lysine respectively). The precursor mass tolerance was set to 7 parts-per-million (ppm) and MS/MS of 10 ppm, with a maximum false discovery rate of 5.0% set for protein identifications and PSM, which was filtered to 1% at the peptide and protein level after peptide score correction. To enhance the identification of peptides between fractions and replicates, the match between runs option was enabled with a precursor match window set to 2 min and an alignment window of 10 min. The mass spectrometry proteomics data have been deposited to the ProteomeXchange Consortium (Vizcaíno et al., 2014) via the PRIDE partner repository (Vizcaíno et al., 2016) with the dataset identifier PXD007288. In addition, processed chromatograms for each tissue have been deposited to the EMBL—EBI BioStudies database (Sarkans et al., 2018), with accession S-BSST152.

### Co-immunoprecipitation

Brain and liver tissues collected from 3 mice (B10, male, 10-week old), were rinsed in ice cold size-exclusion chromatography buffer including protease inhibitors without EDTA (Roche) and phosphatase inhibitors (1 mM sodium orthovanadate, 5 mM sodium pyrovanadate and 0.5 mM pervanadate) and placed on ice. Tissues (100-200 mg) were individually disrupted in a Dounce homogenizer (2 min, tight dounce) in 2 mL of ice-cold buffer. The resulting lysates were clarified by centrifugation (15 min; 4°C; 16,000 r.c.f.) and 1 ml of each supernatant was used for immunoprecipitation using the C-9 mouse monoclonal talin antibody (sc-365875, Santa Cruz) following manufacturer’s protocol. Briefly, the lysates were pre-cleared for 30 min using mouse IgG and protein L-agarose (sc 2336, Santa Cruz), incubated for 30 min with 2 *μ*g of primary antibody, followed by 30 min incubation with protein L-agarose. The bound beads were washed twice with ice-cold SEC buffer and the protein eluted by boiling with the Laemmli buffer. The resulting proteins were subjected to SDS-PAGE, in-gel digested, purified on STAGE tips and analyzed by mass spectrometry. The raw data was searched using Byonic search engine (v.2.7.7, Protein Metrics, San Carlos, CA) and the resulting protein intensity values were used for normalizing protein abundances across different tissues.

### Interactome data analysis

PrInCE (Stacey et al., 2017), an open-source pipeline for co-migration data analysis, was used to reconstruct high-confidence interactomes from PCP-SILAM profiles. PrInCE first performs basic data filtering and preprocessing, then applies a machine-learning approach to rank interactions, and returns as output a list of interactions at a given precision threshold. Briefly, proteins detected in fewer than five fractions are filtered, single missing values are imputed as the mean of neighboring intensities, and co-fractionation profiles are smoothed by a sliding average with a width of five fractions. Next, mixtures of one to five Gaussians are fitted to each profile, and model selection is performed using the bias-corrected Akaike information criterion (Hurvich and Tsai, 1989). Five distance measures are calculated for each protein pair, including the Pearson correlation coefficient and its *P* value, the Euclidean distance, the number of fractions separating the maximum values of each co-fractionation profile, and the Euclidean distance between the closest two Gaussians. A naïve Bayes classifier is subsequently trained on a reference set of interactions in 15-fold cross-validation, using this set of distance measures as features, and protein pairs are ranked as candidate interactions. Here, we used CORUM (Ruepp et al., 2010) (release 30.10.2016) to calculate precision, as well as for all further precision and complex membership analyses. The naïve Bayes classifier calculates an interaction probability for every pair, and this probability is converted to a measure of precision for each interaction by calculating the ratio of true positives to true positives and true negatives among interactions at that probability or higher. Finally, a list of interactions is output at a user-specified precision. We retained interactions within each tissue at 80% precision or higher for further analysis, but additionally provide all interactions at 50% precision in Table S1.

### Comparison of human high-throughput and mouse tissue interactomes

Interaction data from BioPlex (Huttlin et al., 2015), BioPlex 2 (Huttlin et al., 2017), HI-II-14 (Rolland et al., 2014), (Hein et al., 2015), and (Wan et al., 2015) were downloaded from the supporting information of the respective publications. Self-interactions and isoforms were removed, and Ensembl or Entrez identifiers were mapped to UniProt accessions in Bioconductor (Huber et al., 2015). Precision was calculated by defining true positives as intra-complex interactions and false positives as inter-complex interactions (Havugimana et al., 2012). Enrichment for shared Gene Ontology (GO) terms was calculated as the ratio of the proportion of interacting proteins that shared at least one annotation at a given breadth to the proportion of protein pairs within the organism’s proteome sharing at least one annotation at the same breadth (Simonis et al., 2009). Enrichment analysis was performed at three breadths, considering GO terms annotated to fewer than 10, 100, or 1000 proteins. Human and mouse GO annotations were obtained from the UniProt-GOA (Huntley et al., 2015). Annotations with the evidence code “IPI” (inferred from physical interaction) or the qualifier “NOT” were removed. RNA-seq expression data from 77 healthy human samples and 109 healthy mouse samples was obtained from Bgee (Bastian et al., 2008) and used to calculate mRNA coexpression. The Pearson correlation coefficient was used as a measure of coexpression throughout. Known domain-domain interactions were obtained from 3did (Mosca et al., 2014), while Pfam domain annotations were obtained from UniProt (The UniProt Consortium, 2017), and enrichment for known domain-domain interactions was calculated as for shared GO terms.

### Hierarchical clustering of tissue transcriptomes, proteomes, and interactomes

To analyze relationships between tissues on the basis of their interactomes, hierarchical clustering of tissue interactomes was performed using the Jaccard index as the similarity measure. We additionally performed hierarchical clustering using only the induced subgraphs formed by the set of proteins that participated in at least one interaction in each of the seven tissue interactomes, to control for differences at the tissue proteome level. Relationships between tissues were consistent when using all interactions or only interactions made by housekeeping proteins. Tissue transcriptomes from (Merkin et al., 2012) were clustered using the Pearson correlation coefficient as the similarity measure; only the 5,000 most variable genes were used for clustering analysis. Tissue proteomes from (Geiger et al., 2013) were clustered using the Pearson correlation coefficient as the similarity measure.

### Analysis of novel mouse interactions

To analyze mouse interaction novelty, a master database of 72,645 experimentally detected mouse interactions was compiled by merging interactions from BioGrid (Chatr-Aryamontri et al., 2017), CORUM (Ruepp et al., 2010), IID (Kotlyar et al., 2015), IntAct (Orchard et al., 2014), iRefIndex (Razick et al., 2008), mentha (Calderone et al., 2013), and MINT (Licata et al., 2012). Database interactions were mapped to UniProt identifiers using Bioconductor or mapping files provided by the database, and isoforms were removed. The combined set of unique interactions from all seven databases was used to define known interactions and identify interactome orphans as proteins in the mouse tissue interactomes for which no interactions were previously deposited in any interaction database. Proteins of unknown function were defined as proteins without any GO term annotation in the UniProt GOA database. Differences in patterns of co-annotation between LC and PCP-SILAM interactions were quantified as odds ratios, and significance was assessed via a Z test, with Benjamini-Hochberg correction. Markov clustering was performed using the R package MCL, allowing self-loops. The number of publications for each gene was calculated using the NCBI file gene2pubmed. Protein abundance in mouse NIH3T3 mouse fibroblasts was obtained from (Schwanhäusser et al., 2011).

### Comparison to predicted tissue interactomes

Predicted interactomes for six tissues, based on tissue-specific gene expression, were obtained from IID (Kotlyar et al., 2015); a predicted interactome was not available for thymus. Mouse tissue-specific gene coexpression networks were constructed for six tissues; an insufficient number of samples were available for skeletal muscle to construct a reliable network. Microarray samples of healthy mouse tissues from the Affymetrix GeneChip Mouse Genome 430 2.0 platform were identified using Bgee (Bastian et al., 2008) and downloaded from ArrayExpress (Kolesnikov et al., 2015). Samples were processed using BrainArray Custom CDF (Dai et al., 2005) version 21.0.0, mapping probesets to Ensembl gene accessions, and normalized using MAS5 (Hubbell et al., 2002). Probes called as present in fewer than 20% of samples for each tissue were removed. ComBat (Leek et al., 2012) was used to adjust for batch effects, using each experiment as a batch (Vandenbon et al., 2016). Finally, Ensembl gene identifiers were mapped to UniProt accessions using identifier mapping files provided by UniProt.

Enrichment for experimentally detected interactions was calculated as the ratio of overlap between tissue interactomes and tissue-specific coexpression networks and predicted tissue interactomes to the overlap when tissue interactomes were randomly rewired 1,000 times using a degree-preserving algorithm (Maslov and Sneppen, 2002). The number of iterations for the edge rewiring algorithm was set to 6.9 times the number of edges in each network (Ray et al., 2012). Analysis of network topology was performed using the R package igraph (Csardi and Nepusz, 2006). Hub proteins were defined as the top 10% most connected proteins in each network. The tendency for a protein to interact with different partners across tissues was quantified as the mean Jaccard index across all tissue pairs, with a lower Jaccard index reflecting greater rewiring of protein interactions and a higher Jaccard index reflecting relatively stable interactions across tissues. Enrichment or depletion for interactions at each level of specificity was calculated by calculating the ratio of interactions observed at each tissue specificity relative to random expectation, using the same rewired interactomes as above. No interactions were observed in five or more tissues within randomized networks.

### Evolutionary analysis of tissue-specific interactions

Evolutionary conservation of mouse interactions in Saccharomyces cerevisiae, Caenorhabditis elegans, and Drosophila melanogaster was calculated using model organism specific interactions from BioGrid (Chatr-Aryamontri et al., 2017). Mouse proteins were mapped to their one-to-one orthologs in each organism, as well as human, using Ensembl (Kinsella et al., 2011). The difference in evolutionary rates, ΔK, between interacting protein pairs was calculated as in (Fraser et al., 2002). Estimates of protein evolutionary age were obtained from ProteinHistorian (Capra et al., 2012). Phylogenetic profiles were constructed using the InParanoid database (Sonnhammer and Östlund, 2015), with the similarity in phylogenetic profile of a protein pair defined as the Pearson correlation in the presence or absence of each protein across all species (Fortelny et al., 2017). To analyze cross-talk between tissue-specific and housekeeping proteins, we calculated the number of tissues in which each protein was detected in at least one PCP-SILAM fraction. We then calculated the statistical significance of the over or under-representation of interactions between housekeeping proteins and proteins detected in one to seven tissues based on the randomly rewired interactomes described above, using Bonferroni correction to correct for multiple hypothesis testing. To assess statistical significance across all seven tissues, Z scores were aggregated using Stouffer’s method, weighting each Z score by the number of interactions detected in that tissue.

### Analysis of interaction rewiring across tissues

Enrichment analysis of rewired proteins was performed using data from the following sources: proteins involved in stable protein complexes were identified from CORUM (Ruepp et al., 2010). Enrichment for individual functional annotations within the Gene Ontology was evaluated using a hypergeometric test, as implemented in the R package `HTSanalyzeR` (Wang et al., 2011). Protein domains were identified using Pfam version 30.0 (Finn et al., 2016). Intrinsically disordered proteins were identified using IUPred (Dosztányi et al., 2005), with proteins containing more than 30% disordered residues categorized as intrinsically disordered (Gsponer et al., 2008). Proteins containing linear motifs were identified using ANCHOR (Mészáros et al., 2009). The set of mouse disease genes was obtained from the Mouse Genome Database (Eppig et al., 2015). Protein and mRNA abundance, half-lives, and translation and transcription rates were obtained from (Schwanhäusser et al., 2011). Transcription factors were obtained from the Mouse TF Atlas (Zhou et al., 2017). Cell surface protein receptors were identified based on the annotations provided in (Ramilowski et al., 2015). Protein kinases were obtained from the UniProt protein kinase classification and index (https://www.uniprot.org/docs/pkinfam). Edge betweenness centrality was calculated in the aggregate PCP—SILAM interactome formed by the union of unique interactions across all seven tissues, using the R package igraph (Csardi and Nepusz, 2006).

---

**Figure S1.**
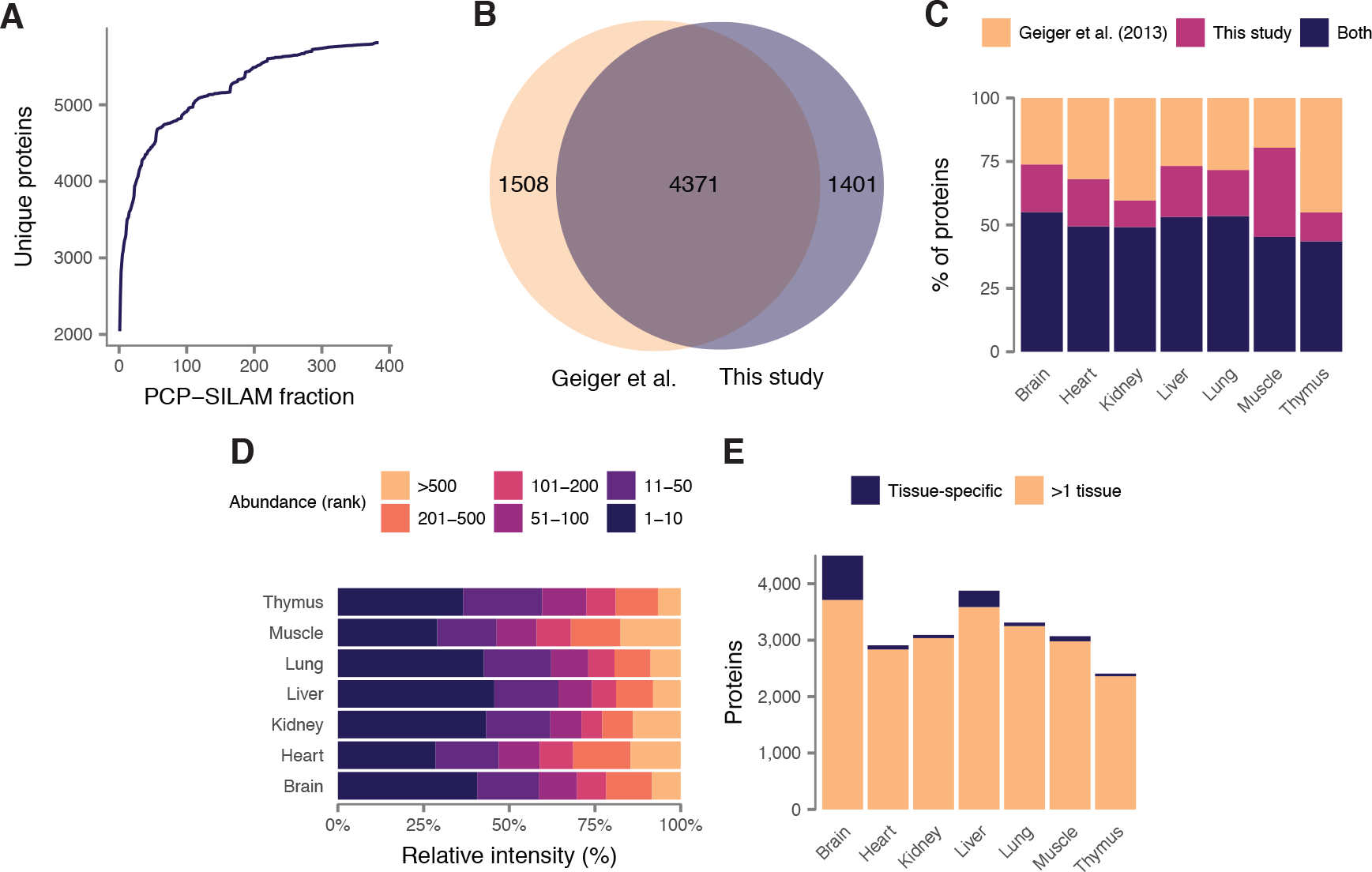
Quantitative profiling of mouse tissue proteomes by PCP—SILAM. Related to Figure 1. (A) Saturation of unique proteins identified with additional PCP fractions, consistent with the presence of the SILAM global reference. (B) Unique proteins identified across all seven mouse tissues by this study and (Geiger et al., 2013). (C) Unique proteins identified within each mouse tissue by this study and (Geiger et al., 2013). (D) Distribution of protein abundances within tissue interactomes. Summed ion intensities of ranked proteins are presented as a percentage of the total summed ion intensities. (E)(Proteins identified only in a single tissue (tissue-specific), or in at least one other tissue, within each mouse tissue proteome.)

**Figure S2.**
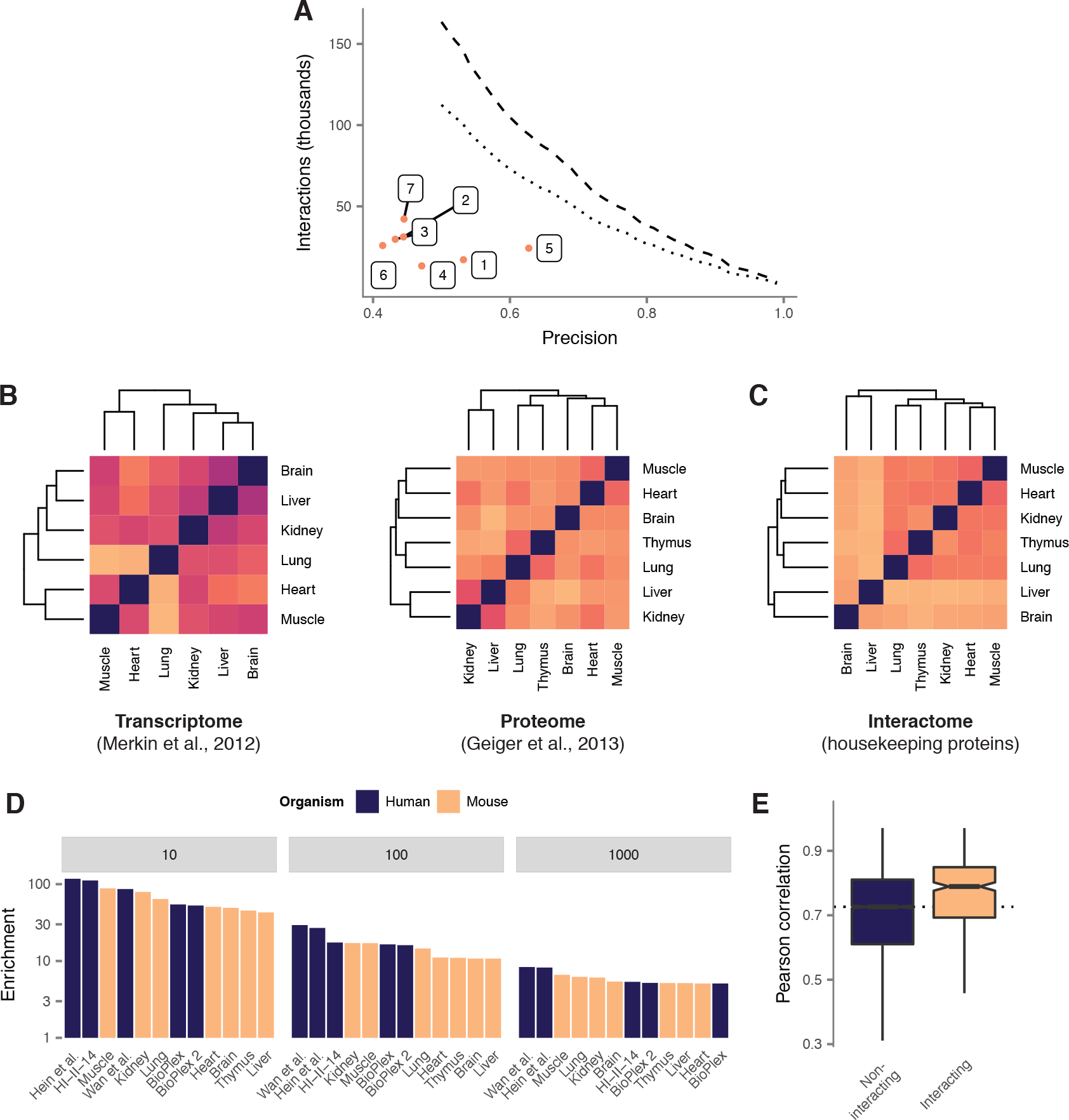
Bioinformatic validation of PCP-SILAM mouse tissue interactomes. Related to Figure 2. (A) Precision of mouse tissue interactomes compared to seven mouse literature-curated protein-protein interaction databases. Dashed line shows all interactions; dotted line shows unique interactions. 1, BioGrid (Chatr-Aryamontri et al., 2017); 2, HINT (Das and Yu, 2012); 3, IID (Kotlyar et al., 2015); 4, IntAct (Orchard et al., 2014); 5, iRefIndex (Razick et al., 2008); 6, mentha (Calderone et al., 2013); 7, all unique interactions from mouse interaction databases 1-6. (B) Hierarchical clustering of mouse tissue transcriptomes (Merkin et al., 2012) and proteomes (Geiger et al., 2013). (C) Hierarchical clustering of mouse tissue interactomes based on induced subgraphs formed by proteins that participated in at least one interaction at 80% precision in each of the seven tissues (“housekeeping” proteins). (D) Enrichment for shared Gene Ontology (GO) terms between interacting protein pairs in mouse tissue interactomes and five recent high-throughput human interactome maps, considering only GO terms annotated to fewer than 10, 100, or 1000 proteins within the human and mouse proteomes. (E)Pearson correlations across huntingtin affinity purifications in mouse brains (Shirasaki et al., 2012) between interacting and non-interacting protein pairs in mouse brain, as resolved by PCP-SILAM.

**Figure S3.**
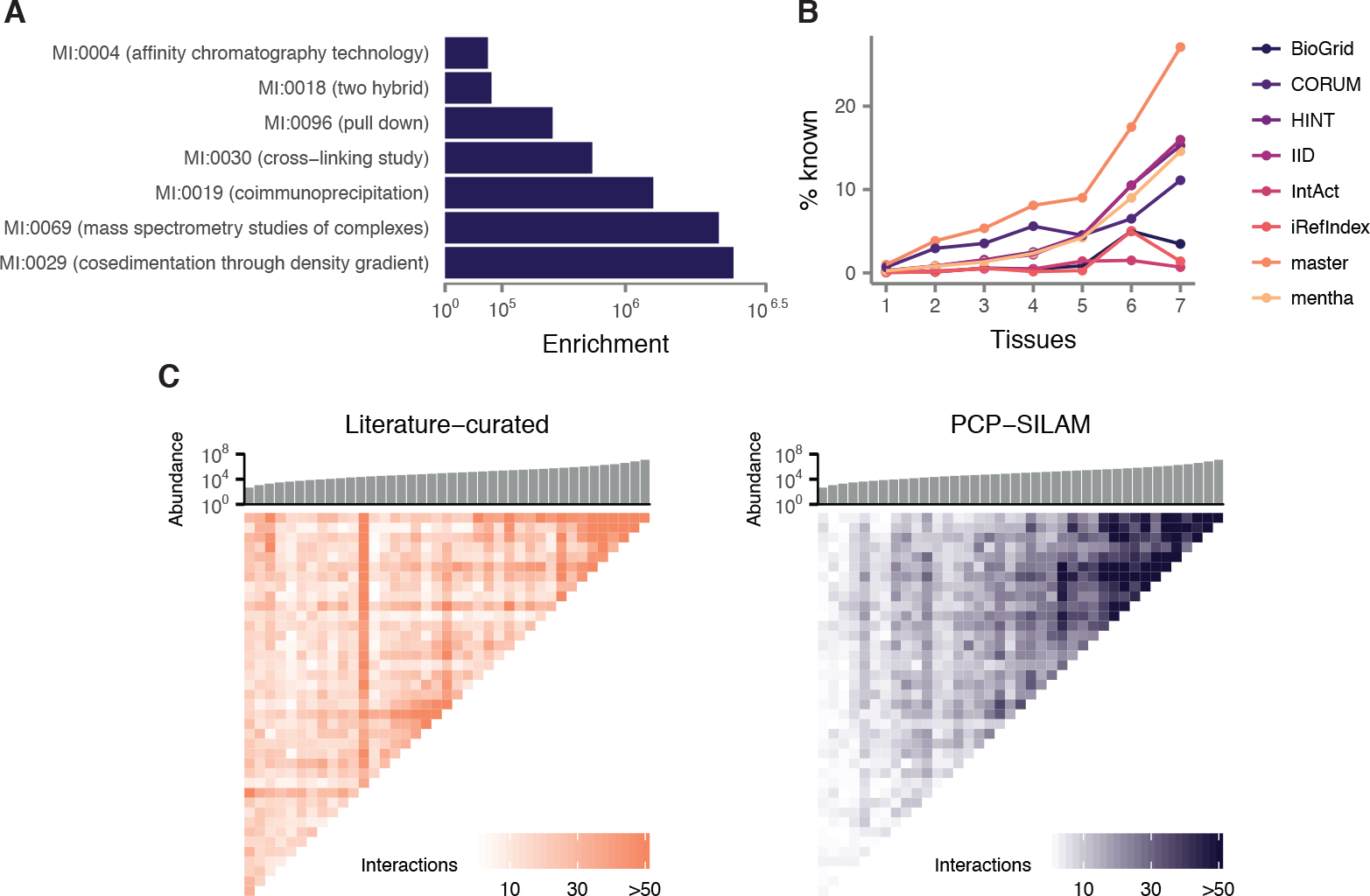
Expansion of the known mouse interactome by PCP-SILAM. Related to Figure 3. (A) Enrichment for overlap with literature-curated mouse interactions detected with different methods, relative to random expectation. (B) Proportion of previously known interactions from seven literature-curated mouse interaction databases, and a master list of all known mouse interactions, for PCP-SILAM interactions detected in one to seven tissues. (C) Number of interactions between proteins binned by mean abundance (copies per cell) in mouse fibroblasts (Schwan-hausser et al., 2011), and ordered along both axes. Histogram shows the median of abundance in each bin.

**Figure S4.**
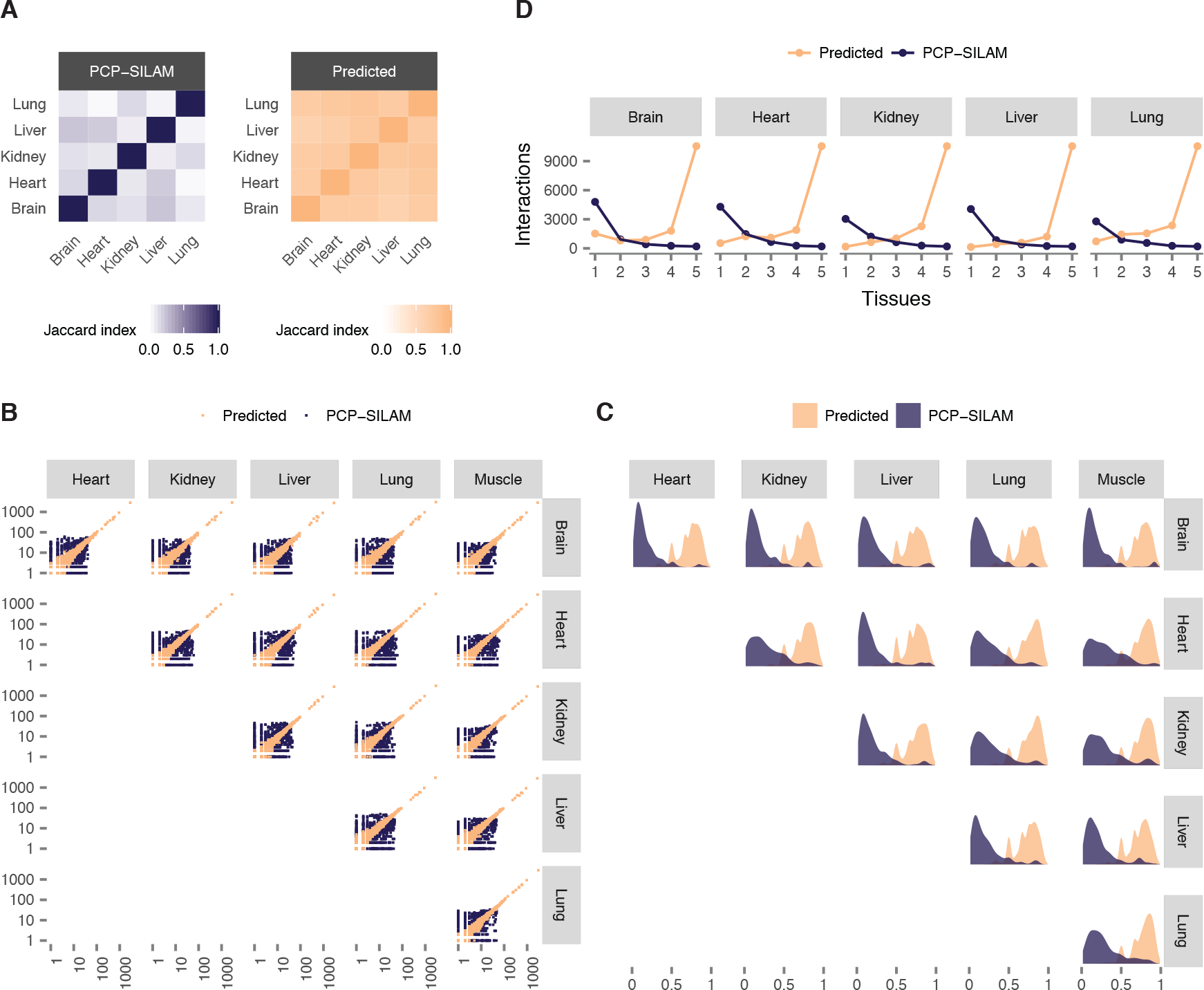
Interactome rewiring limits acuracy of tissue interactome prediction. Related to Figure 4. (A) Overlap in hub proteins between predicted and PCP-SILAM tissue interactomes. (B) Protein degrees across tissues in predicted and PCP-SILAM tissue interactomes. (C) Rewiring of protein interaction partners across tissues (Jaccard index) in predicted and PCP-SILAM tissue interactomes. (D) Tissue specificity of interactions across predicted and PCP—SILAM tissue interactomes.

**Figure S5.**
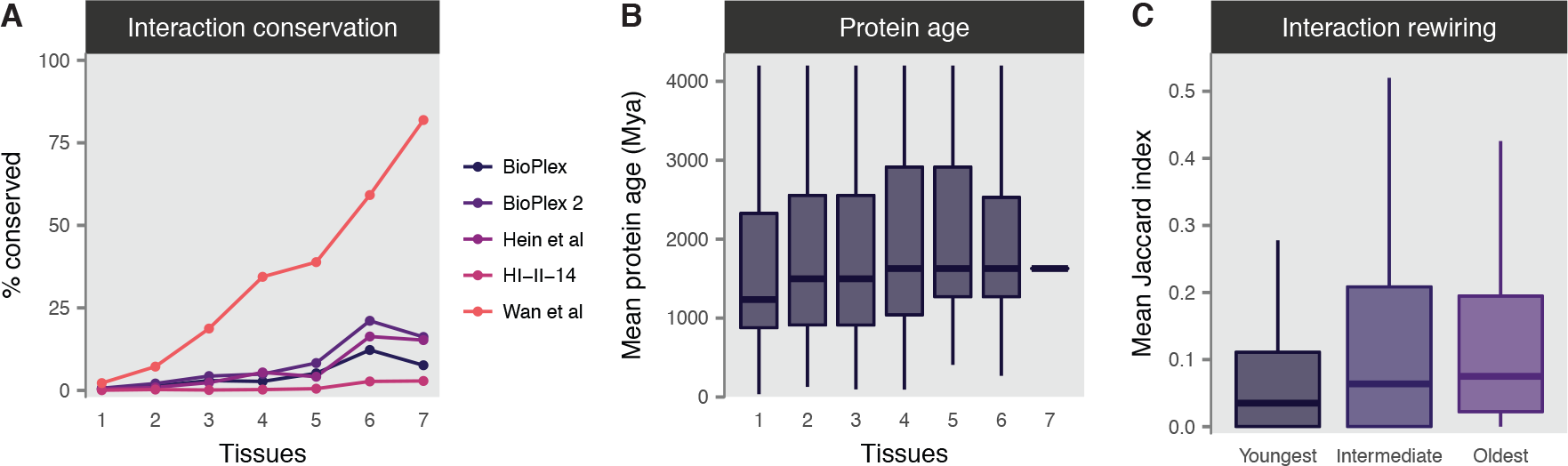
Evolution of interactions in mouse tissues. Related to Figure 5. (A) Proportion of mouse interactions conserved in human at each level of tissue specificity, using five recent high throughput human interactome screens. (B) Mean evolutionary age of interacting protein pairs at each level of tissue specificity. (C) Mean Jaccard index across all pairs of tissue interactomes for mouse proteins binned into terciles by evolutionary age.

**Figure S6.**
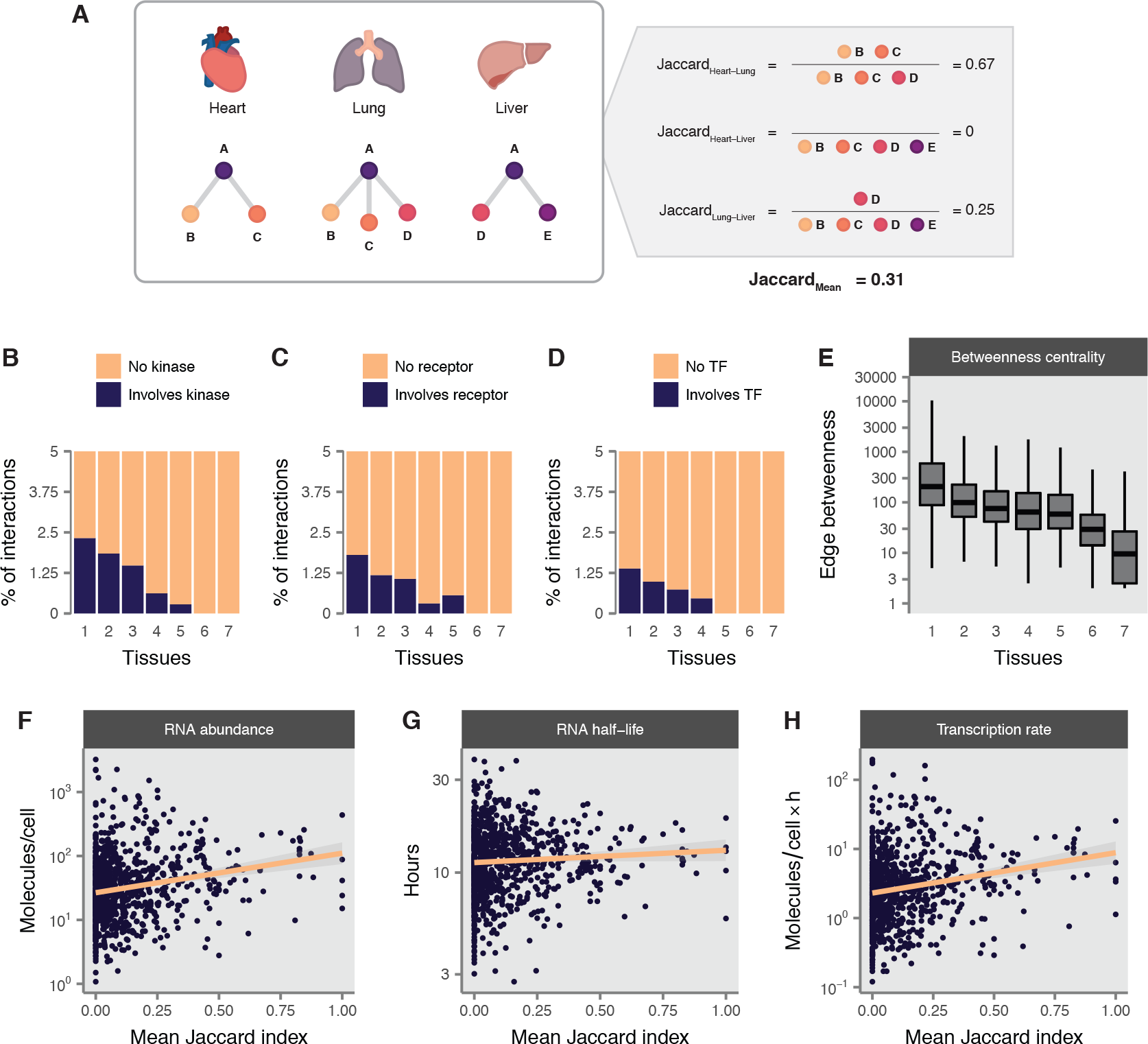
Tissue-specific interactions mediate tissue-specific biological information flow. Related to Figure 6. (A) Overview of the mean Jaccard index calculation for protein A with interaction partners B, C, D, and E in three tissues. (B) Proportion of interactions involving protein kinases at each level of tissue specificity. (C) Proportion of interactions involving transcription factors at each level of tissue specificity. (D) Proportion of interactions involving cell surface receptors at each level of tissue specificity. (E) Betweenness centrality of interactions at each level of tissue specificity. (F-H) mRNAs encoding rewired proteins are characterized by low abundance (F), short half-lives (G), and slow transcription rates (H).

## Supplementary tables

**Table S1**

PCP-SILAM interactomes of seven mouse tissues at 50% precision.

**Table S2**

Compilation of known mouse interactions from seven literature-curated protein interaction databases.

**Table S3**

(A) Proteins of unknown function in mouse interactomes and their interacting partners in each tissue interactome.

Interactome orphans in mouse interactomes and their interacting partners in each tissue interactome.

